# Towards Effective and Generalizable Fine-tuning for Pre-trained Molecular Graph Models

**DOI:** 10.1101/2022.02.03.479055

**Authors:** Jun Xia, Jiangbin Zheng, Cheng Tan, Ge Wang, Stan Z. Li

## Abstract

Graph Neural Networks (GNNs) and Transformer have emerged as dominant tools for AI-driven drug discovery. Many state-of-the-art methods first pre-train GNNs or the hybrid of GNNs and Transformer on a large molecular database and then fine-tune on downstream tasks. However, different from other domains such as computer vision (CV) or natural language processing (NLP), getting labels for molecular data of downstream tasks often requires resource-intensive wet-lab experiments. Besides, the pre-trained models are often of extremely high complexity with huge parameters. These often cause the fine-tuned model to over-fit the training data of downstream tasks and significantly deteriorate the performance. To alleviate these critical yet under-explored issues, we propose two straightforward yet effective strategies to attain better generalization performance: 1. MolAug, which enriches the molecular datasets of down-stream tasks with chemical homologies and enantiomers; 2. WordReg, which controls the complexity of the pre-trained models with a smoothness-inducing regularization built on dropout. Extensive experiments demonstrate that our proposed strategies achieve notable and consistent improvements over vanilla fine-tuning and yield multiple state-of-the-art results. Also, these strategies are model-agnostic and readily pluggable into fine-tuning of various pre-trained molecular graph models. We will release the code and the fine-tuned models.

## 1 Introduction

Pre-trained language models (PLMs) have foundamentally changed the landscape of natural language processing (NLP) [Devlin *et al*., 2019], which have established new state-of-the-art results for a large variety of NLP tasks. Inspired by their proliferation, tremendous efforts have been devoted to molecular graph pre-training which can exploit abundant knowledge of unlabelled molecular in the database [Hu *et al*., 2020]. For the pre-training stage, existing works train the encoder with various pretext tasks in absence of labels [Rong *et al*., 2020]. For the second fine-tuning stage, researchers adapt the pre-trained models to the downstream tasks via replacing the top layer of the pre-trained models by a task specific sub-network, and then continuing to train the new model with the limited data of the downstream task.

Despite the fruitful progress in the strategies for pre-training, the fine-tuning stage remains under-explored for pre-trained molecular graph models. There are two crucial issues impede the improvements of performance during fine-tuning: (1) insufficient labeled data for downstream tasks. Different from other domains that have abundant labeled data, getting high-quality labels for molecular data often requires resource-intensive wet-lab experiments [Xia *et al*., 2021a]. (2) the pre-trained models are often of extremely high complexity with tens millions of parameters [Rong *et al*., 2020; Li *et al*., 2021b], which posses the capability of memorizing the limited samples and lead to poor generalization [Mohri *et al*., 2012; Haoming *et al*., 2020]. As shown in Figure 1, we conduct vanilla fine-tuning on one of the state-of-the-art pre-trained molecular models MPG [Li *et al*., 2021b]. The over-fitting issue poses hurdle to the further improvements on various datasets. To mitigate this issue, existing fine-tuning methods in other domains often rely on hyper-parameter tuning heuristics. For example, Howard and Ruder [Howard and Ruder, 2018] follow a heuristic learning rate schedule and gradually unfreeze the layers of the pre-trained language model to improve the fine-tuning performance, which require significant tuning efforts. The other line of works propose various regularizations to control complexity of pre-trained models. Specifically, SMART [Haoming *et al*., 2020] introduces a regularization, which encourages the output of the model not to change much, when injecting an adversarial perturbation to the input. However, this strategy is not suitable for pre-trained graph models because synthesizing adversarial samples for molecular graphs online is time-consuming for its high complexity. Recently, Child-Tuning [Xu *et al*., 2021b] is proposed to strategically mask out some gradients of network during the backward process. Albeit effective, it suffers a heavy computational overhead because it requires to search for suitable masks for each parameter in the pre-trained models. Now, we are naturally motivated to ask following question: *Can we devise more suitable strategies for effective and generalizable fine-tuning on pre-trained molecular graph models?*

**Figure 1:**
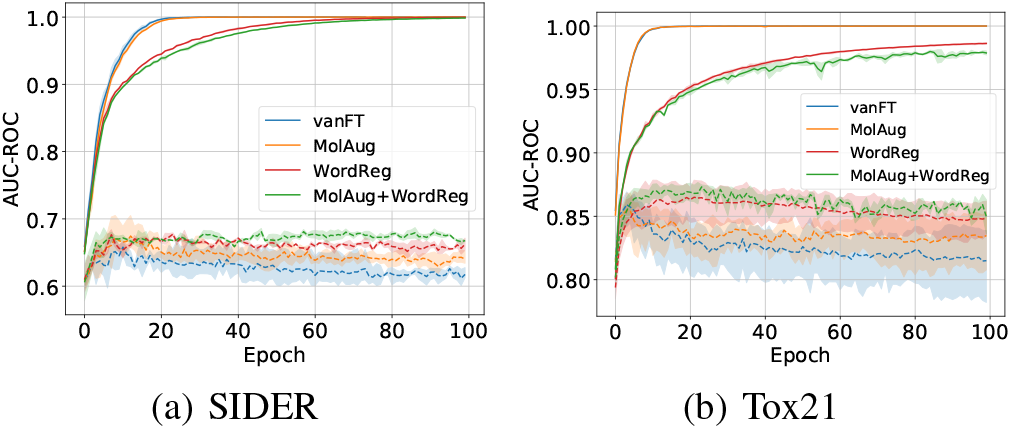
Training (solid lines) and testing (dashed lines) curves of various fine-tuning strategies on SIDER and Tox21 datasets. ‘vanFT’ refers to vanilla fine-tuning. The over-fitting issue of vanilla fine-tuning impede the performance improvements while our proposed MolAug, WordReg or their combination can alleviate this issue.

To fully harness the power of pre-trained molecular graph models, we propose two strategies for better fine-tuning: MolAug and WordReg. MolAug is molecular graph augmentation with chemical enantiomers and homologies which share the similar (or identical) physical (permeability, solubility, etc.) or chemical (toxicity, etc.) properties with themselves. WordReg is a novel smoothness-inducing regularization built on dropout. To the best of our knowledge, we are the first to study the fine-tuning stage of the pre-trained molecular graph models, which is important while being neglected. We summarize our contributions here:

- We propose new data augmentations for molecular graph data, which introduce variations while not altering the physical or chemical properties of molecules too much.
- We propose a novel smoothness-inducing regularization built on dropout that can effectively control the high complexity of the pre-trained models. Also, it is domain-agnostic and we study its effects in fine-tuning pre-trained models of NLP and CV in the appendix.
- Through extensive experiments on 3 drug discovery sub-tasks (11 datasets in total) including molecular property prediction, drug-drug interaction prediction and drug-target interaction prediction, we observe consistent and notable improvements over vanilla fine-tuning and yield multiple new state-of-the-art results.

## 2 Related work

### 2.1 Molecular Representation Learning

Molecular Representation Learning refers to represent molecules in the vector space. Initially, the traditional chemical fingerprints such as ECFP [Rogers and Hahn, 2010] encode the neighbors of atoms in the molecule into a fix-length vector. With the proliferation of deep learning, [Duvenaud *et al*., 2015] first introduce convolutional neural networks to learn neural fingerprints of molecules. Subsequently, some researchers feed the SMILES (a line notation for describing the structure of chemical species using short ASCII strings) into RNN-based models to obtain molecular representations. In order to fulfill the topology information of molecular graphs, some recent works [Kearnes *et al*., 2016; Xiong *et al*., 2020] attempt to apply GNNs to molecular representation learning. Besides, MPNN [Gilmer and others., 2017] and its variants DMPNN [Yang *et al*., 2019], CMPNN [Song *et al*., 2020], CoMPT [Chen *et al*., 2021] utilize a message passing framework to better capture the interactions among atoms. However, they still require expensive annotations and barely generalize to unseen molecules, which pose hurdle to the practical applications.

### 2.2 Pre-training on Graphs

To address the fundamental challenges of extremely scarce labeled data and out-of-distribution generalization in graphs learning, tremendous efforts have been devoted to pre-training on graphs recently. One line of these works follow the contrastive paradigm [Zhu *et al*., 2021b; Zhu *et al*., 2021a; Xia *et al*., 2021b]. For molecular pre-training, GraphCL [You *et al*., 2020] and its variants [You *et al*., 2021; Susheel *et al*., 2021; Xia *et al*., 2022] embeds augmented versions of the anchor molecular graph close to each other and pushes the embeddings of other molecules apart. The other line of works adopt generative pretext tasks. Prototypical examples are GPT-GNN [Hu and others., 2020] which introduces a self-supervised attributed graph generation task to pre-train GNNs so that they can capture the structural and semantic properties of the graph. For molecular graph pre-training, Hu et al. [Hu *et al*., 2020] conduct attribute and structure prediction at the level of individual nodes as well as entire graphs. To capture the rich information in molecular graph motifs, GROVER [Rong *et al*., 2020] and MGSSL [Zhang *et al*., 2021] propose to predict or generate the motifs. Analogously, MPG [Li *et al*., 2021b] learns to compare two half-graphs (each decomposed from a graph sample) and discriminate whether they come from the same source. Despite the progress in molecular graph pre-training, few efforts have been devoted to the fine-tuning except for a recent work [Han and others., 2021] that adaptively selects and combines various auxiliary tasks with the target task in the fine-tuning stage to improve performance, which is impractical because auxiliary tasks are often unavailable during fine-tuning.

## 3 Methodology

### 3.1 MolAug: <u>Mol</u>ecular Graph <u>Aug</u>mentations with Chemical Enantiomers and Homologies

Initially, GraphCL [You *et al*., 2020] augments molecular graph data in the form of naive random corruption (e.g., dropping bonds, dropping atoms and etc.). However, these augmentations may alter molecular graph semantics completely even if the perturbation is weak. For example, dropping a carbon atom in the phenyl ring will alter the aromatic system and result in an alkene chain, which will drastically change the molecular properties. Besides, MoCL [Sun *et al*., 2021] attempts to incorporate domain knowledge into graph data augmentation via replacing valid substructures in molecular graph with bioisosteres that share similar properties. However, bioisosteres are used to modify some molecular properties as expected (e.g, reduce toxicity) in drug design [Mannhold *et al*., 2012], which may introduce incorrect supervision for molecular properties prediction such as toxicity prediction. We compare MolAug with these augmentations in section 4.5.

To alleviate these issues, we resort to chemical enantiomers and homologies. As shown in Figure 2 (a) and (b), an enantiomer is one of two stereoisomers that are mirror images of each other that are non-superposable, much as one’s left and right hands are mirror images of each other that cannot appear identical simply by reorientation. Despite the structural difference, enantiomers share the identical chemical and physical properties with each other in most cases and thus being a more suitable alternative for molecular graph augmentations in molecular properties prediction. In practice, we can obtain chemical enantiomers of a molecule by changing the chirality type which is provided in the atom (node) feature. Besides, as shown in Figure 2 (c) and (d), chemical homologies are a series of compounds differing from each other by a repeating unit, such as a methylene bridge, which share the same chemical properties with each other. Therefore, chemical homologies can serve as an ideal way of augmentation when predicting molecular chemical properties. For each molecule **x** and its label *y* in the dataset 𝒟 of down-stream task, we can obtain its various augmented versions with operation MolAug(·) which can be enantiomers, homologies or their combinations. Now, we can formulate the loss of fine-tuning with MolAug as,

**Figure 2:**
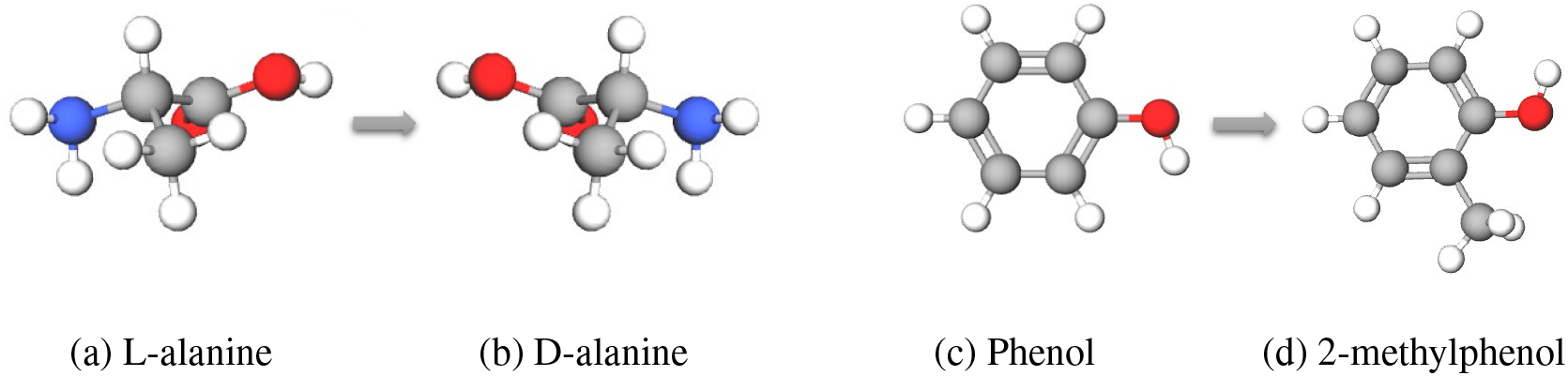
Illustration of MolAug. L-alanine (a) and D-alanine (b) is a pair of enantiomers; Phenol and 2-methylphenol is a pair of homologies. Gray, red, blue and white balls denote Carbon (C), Oxygen (O), Nitrogen (N) and Hydrogen (H) atoms respectively.

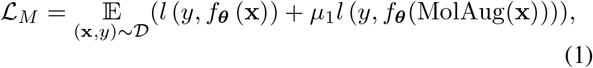

where *μ*_1_ is a trade-off parameter controlling the impact of MolAug and *l* is the loss function of downstream task. We can also obtain various augmentations for each molecule during each iterations with MolAug, which we study in section 4.6. Apart from pretrain-then-finetune paradigm, MolAug can also work well in training from scratch, which we validate in the appendix.

### 3.2 WordReg: Smoothness-inducing Regularization Built on Dropout

Existing fine-tuning strategies [Haoming *et al*., 2020; Xu *et al*., 2021b; Aghajanyan *et al*., 2021] in NLP enforce the output of the model not to change much, when injecting perturbations to the input or model parameters. In other words, they encourage the model to be insensitive to perturbations, and thus effectively control its capacity [Mohri *et al*., 2012]. Compared with these strategies, WordReg imposes the perturbations on the neurons which is more efficient than perturbing the input or huge model parameters. Besides, as shown in Figure 3, WordReg obtains the worst-case dropout via maximizing the divergence between the outputs with two different dropout which is task-dependent and can introduce stronger regularization than random dropout (validated in Figure 6). Formally, consider the pre-trained neural networks *f*_*θ*_(**x, *m***) that takes sample **x** as input when fine-tuning on downstream task. ***m*** is a mask vector denoting which neuron of the pre-trained model should be dropped. More specifically, for the *i*-th unit *m*_*i*_ of vector ***m***, *m*_*i*_ = 1 indicates that the neuron should be dropped while *m*_*i*_ = 0 illustrates that the neuron should be preserved during dropout. To start, we introduce ***m***_*r*_ to denote the mask vector of random dropout of vanilla training. With the dropout ratio *σ* ∈ [0, 1] preseted, we can define the constraint on ***m*** as,

**Figure 3:**
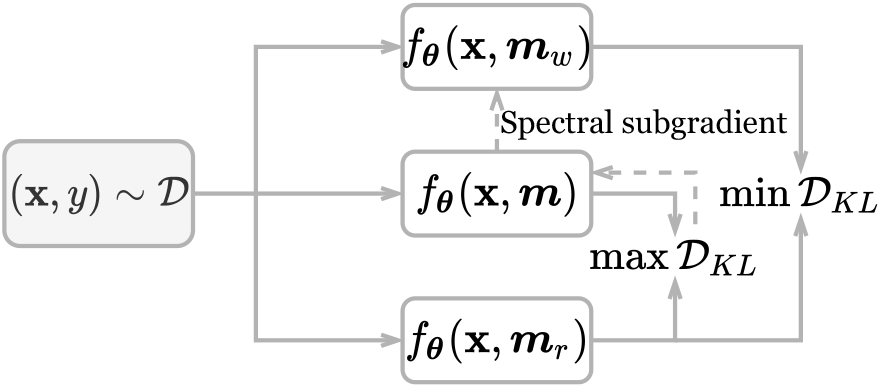
Illustration of WordReg. We obtain the worst-case drop mask vector ***m***_*w*_ (corresponding to ***m***_*r*_) via solving a BQP problem with spectral subgradient-based method.

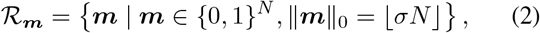

where ∥ · ∥_0_ is the *ℓ*_0_ norm, *N* is the number of neurons in the model. Therefore, we can obtain the worst-case mask vector ***m***_*w*_ by solving an optimization problem,

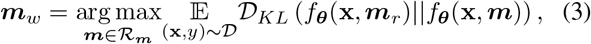

where 𝒟_*KL*_ is Kullback–Leibler divergence. With Taylor expansion, we can approximate the optimal solution as,

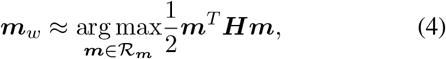

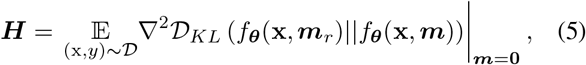

***H*** is the Hessian matrix of the loss 𝒟_*KL*_ at ***m*** = **0** which is a *N* × *N* semi-positive definite matrix. Obviously, this is a Binary Quadratic Programming (BQP) problem, which is NP-hard but admits an approximate solution to ***m***_*w*_. Here, we propose a novel method to solve this problem efficiently based on the spectral subgradient [Boyd *et al*., 2003]. Firstly, we convert {0, 1}-constraint on ***m*** of ℛ_***m***_ to {−1, 1}-constraint on ***n*** via defining ***n*** = 2***m*** − 1. Then we introduce a new variable 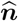 with *N* + 1 dimensions,

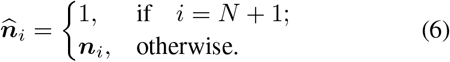

Then we can rewrite constraint ℛ_***m***_ as a new constraint 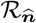 on 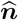,

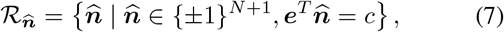

where *c* = 2 ⌊*σN*⌋−*N* +1 and ***e*** ∈ ℝ^*N* +1^ is an all-one vector. By these transformations, we can reformulate the BQP in Eq (5) as a new BQP in terms of 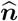, where the constraint term 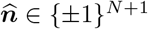 can be rewrited as 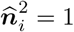. We then introduce a Lagrange multiplier *λ*_*i*_ for each constraint 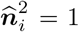 and *λ*_0_ for the constraint 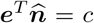. Now, we can formulate the dual problem of the original BQP as,

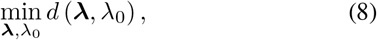

with

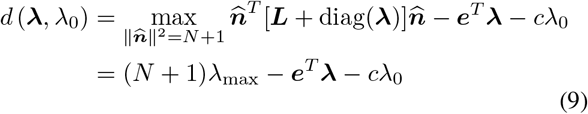

where

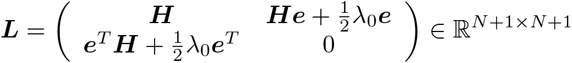

and *λ*_max_ is the largest eigenvalue of ***L*** + diag(***λ***). The eigenvector of unit norm **u**_max_ corresponding to *λ*_max_ can be derived via approximated by using a single-step power iteration instead of conducting naive eigenvalue decomposition. Then, the maximum 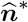 can be derived,

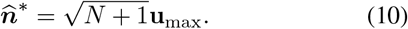

The dual problem Eq.(8) can be solved by the gradient descent method over iterations. We show the gradient of *d* w.r.t ***λ*** and *λ*_0_ as follows,

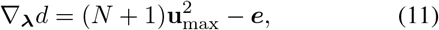

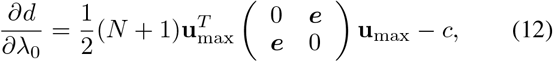

where 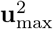 denotes an element-wise square of **u**_max_. During fine-tuning the pre-trained models with WordReg, over each mini-batch, we compute the above gradient to make an one-step update of the Lagrange multipliers ***λ*** and *λ*_0_ along the descending direction, before the maximum 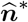 is taken with the updated multipliers. Finally, both 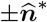 are optimal and we should choose the one closer to 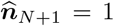 as required. Note that we seek the worst-case dropout mask layer-by-layer instead of applying it to an entire network as a whole. This can make the WordReg computationally efficient as well as prevent too many neurons from being dropped at a few layers. With the worst-case dropout mask vector ***m***_*w*_ obtained, we can formulate the loss of fine-tuning with WordReg as,

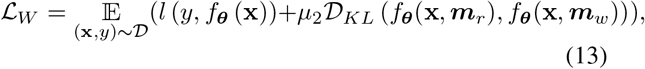

where *μ*_2_ is a trade-off parameter controlling the impact of WordReg, which we study in section 4.6.

#### Remark

There are also some regularizations ELD [Ma *et al*., 2017] and FD [Zolna *et al*., 2018] built on dropout. WordReg differs from them in two aspects: (1) WordReg introduces the worst-cast dropout, which is task-dependent and possess stronger regularization ability than their random drop; (2) ELD and FD are designed to reducing the gap between training (sub-model with dropout) and inference (full model without dropout) while WordReg controls the gap between sub-model with different dropouts.

### 3.3 Combining MolAug and WordReg

Additionally, we can conduct fine-tuning with MolAug and WordReg simultaneously. Formally, we specify the loss function of combining MolAug and WordReg in Eq (14).

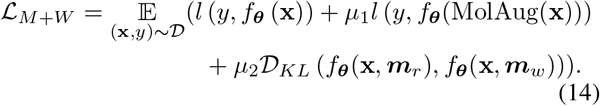

The pseudo codes can be found in the appendix.

## 4 Experiments

### 4.1 Experimental Setup

#### Base Pre-trained Molecular Graph Models

We adopt recent proposed pre-trained molecular graph models MPG [Li *et al*., 2021b] as the base model and fine-tune it with our proposed strategies. We also adopt MGSSL [Zhang *et al*., 2021] and GraphLog [Xu *et al*., 2021a] as our base models to valuate that our strategies can be plugged into the fine-tuning of various pre-trained molecular graph models.

#### Fine-tuning Tasks and Datasets

Following previous works, we use 8 benchmark datasets from the MoleculeNet [Wu *et al*., 2018] to perform molecular property prediction. To keep fair, we adopt the scaffold splitting method with a ratio for train/validation/test as 8:1:1 as previous works. We apply a grid search procedure to obtain the best hyper-parameters with validation set. More details of the hyper-parameters and datasets are deferred to the appendix. *The experiments on drug-drug interaction prediction and drug-target interaction prediction can be found in the appendix*.

#### Baselines

We adopt two types of baselines:

- *Training from scratch:* TF_Roubust [Rogers and Hahn, 2010] is a DNN-based multitask framework taking the molecular fingerprints as the input. GCN [Kipf. and Welling, 2017], Weave [Kearnes *et al*., 2016] and SchNet [Schuett *et al*., 2017] are three graph convolutional models. MPNN and its variants DMPNN, MGCN, CMPNN and CoMPT have been introduced in the related work. AttentiveFP [Xiong *et al*., 2020] is an extension of the graph attention network.
- *Pretrain-then-finetune:* We also compare with several pre-trained models with vanilla fine-tuning: Mol2Vec [Jaeger *et al*., 2018], N-Gram [Liu *et al*., 2019], SMILES-BERT [Wang *et al*., 2019], GROVER, Hu et.al, MPG and several graph contrastive learning methods as introduced in the related work.

### 4.2 Comparisons with State-of-the-art Results

We first fine-tune the state-of-the-art pre-trained molecular graph model MPG with MolAug, WordReg and their combinations respectively. The results are reported in Table 1 which provides the following observations: (1) MolAug and WordReg consistently perform better than vanilla fine-tuning on MPG. The overall relative improvement is 7.2% (3.4% on classification tasks and 18.4% on regression tasks); (2) Equipped with MolAug, WordReg or their combinations, MPG yields new state-of-the-art results on molecular property prediction; (3) The combination of MolAug and WordReg brings a 26.4% relative improvement over MPG on a small dataset FreeSolv with only 642 labeled molecules. This confirms that our strategies can alleviate the over-fitting issues which is often severer in small-scaled datasets.

**Table 1:**
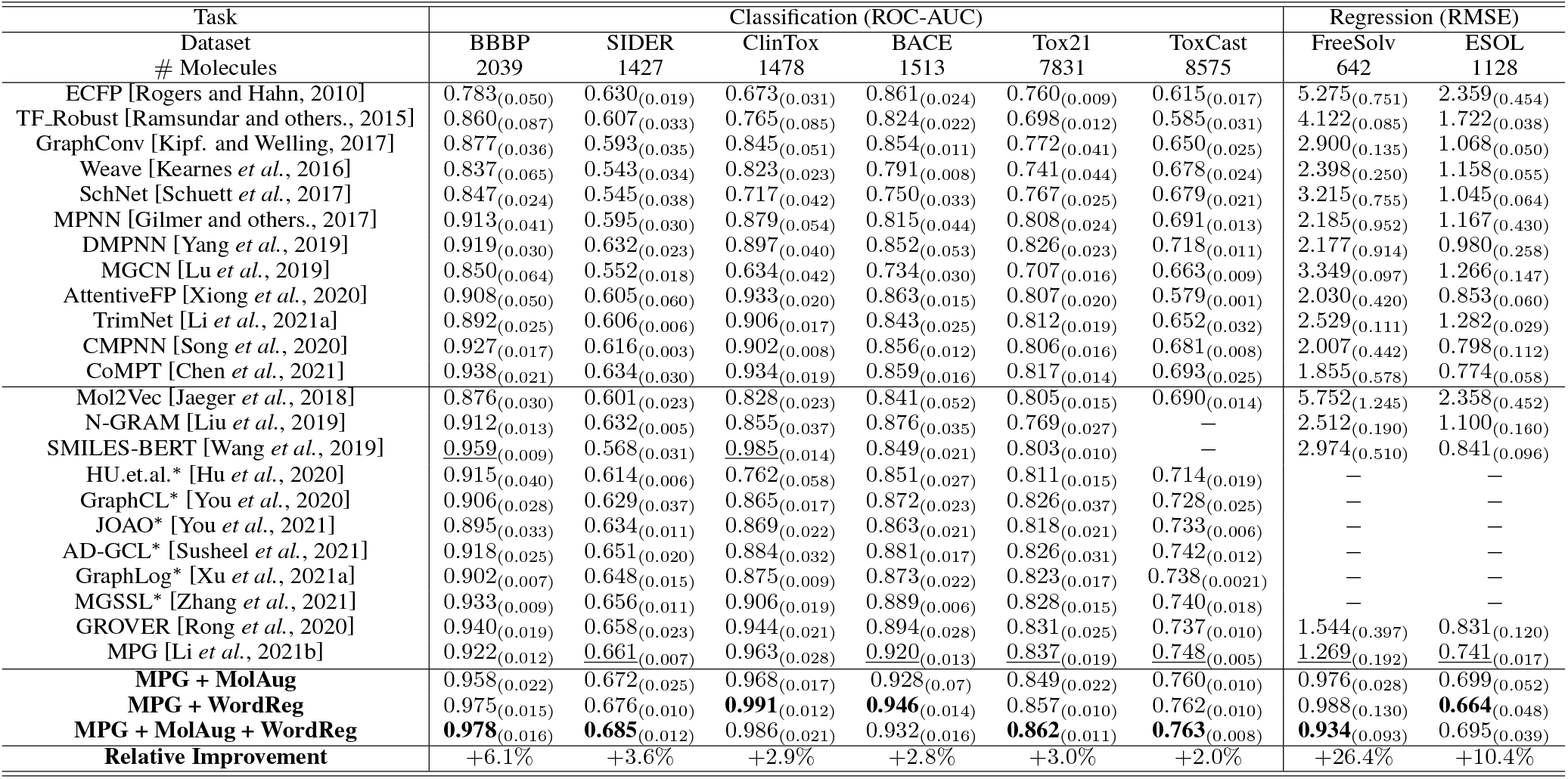
The performance comparison (*higher is better for classification task, lower is better for regression task*). *Relative Improvement* refers to the absolute improvement over MPG base model divided by the original results of MPG. We adopt it as a unified description of improvements for both classification and regression. The methods marked with ‘*’ means the original papers follow a different data splitting with MPG, we fine-tune their public pre-trained models with the splitting as MPG and report the results here. The numbers in brackets are the standard deviation and the ones underlined are the previous best results. ‘−’ means the methods are too time-consuming or original implementations do not admit regression task without non-trivial modifications.

### 4.3 Comparisons with Other Fine-tuning Methods

Due to the fact that there is no fine-tuning methods specially designed for pre-trained molecular graph models, we consider some start-of-the-arts fine-tuning strategies in other domains. As can be observed in Table 2, generally, existing fine-tuning techniques can also bring improvements for Pre-trained molecular graph models. Compared with them, MolAug and WordReg are more suitable for pre-trained molecular graph models because they can avoid heavy tunning or heavy computational overhead of other fine-tuning methods and achieve the best performance among them.

**Table 2:** Comparison with other fine-tuning methods. The running time is evaluated on the same device (Tesla V100 GPU). We provide more details and results in the appendix.

| Methods | BBBP |  | SIDER |  |
| --- | --- | --- | --- | --- |
|  | ROC-AUC | Time (s) | ROC-AUC | Time (s) |
| Vanilla Fine-tuning (Base model) | 0.922<sub>(0.012)</sub> | 654.6<sub>(7.9)</sub> | 0.661<sub>(0.007)</sub> | 327.6<sub>(5.2)</sub> |
| SMART [Haoming et al., 2020] | 0.959<sub>(0.009)</sub> | 9583.3<sub>(26.2)</sub> | 0.673<sub>(0.005)</sub> | 8927.5<sub>(31.1)</sub> |
| R3F [Aghajanyan et al., 2021] | 0.931<sub>(0.008)</sub> | 3876.5<sub>(35.6)</sub> | 0.659<sub>(0.013)</sub> | 2930.5<sub>(16.7)</sub> |
| Adaptive Fine-tuning [Han and others., 2021] | 0.944<sub>(0.012)</sub> | 8051.2<sub>(8.2)</sub> | 0.653<sub>(0.012)</sub> | 6982.3<sub>(15.9)</sub> |
| ChildTuning [Xu et al., 2021b] | 0.952<sub>(0.005)</sub> | 8541.4<sub>(24.8)</sub> | 0.668<sub>(0.017)</sub> | 7358.6<sub>(22.8)</sub> |
| MolAug | 0.958<sub>(0.027)</sub> | 789.8<sub>(15.1)</sub> | 0.672<sub>(0.019)</sub> | 405.6<sub>(5.7)</sub> |
| WordReg | <u>0.975</u><sub>(0.015)</sub> | 1659.8<sub>(10.1)</sub> | <u>0.676</u><sub>(0.010)</sub> | 1436.3<sub>(20.3)</sub> |
| MolAug + WordReg | **0.978**<sub>(0.016)</sub> | 2322.4<sub>(16.8)</sub> | **0.685**<sub>(0.012)</sub> | 1791.68<sub>(14.9)</sub> |

### 4.4 Improving the Performance of Various Pre-trained Molecular Graph Models

To validate that MolAug and WordReg are model-agnostic, we fine-tune other pre-trained molecular graph models including MGSSL and GraphLog (see in the appendix for the limited space) with our MolAug and WordReg. Figure 4 shows the comparisons between our strategies and vanilla fine-tuning on the pre-trained model MGSSL. Obviously, our strategies outperform vanilla fine-tuning across various datasets, which validates that MolAug & WordReg can also improve the performance when fine-tuning various pre-trained molecular graph models.

**Figure 4:**
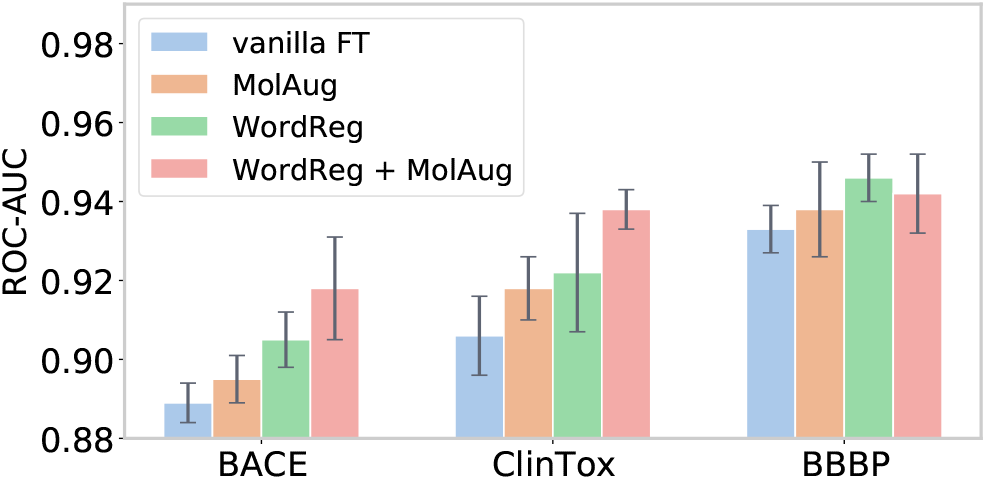
MolAug and WordReg’s performance (compared with vanilla fine-tuning) on another pre-trained model MGSSL.

### 4.5 Ablation Study

#### Ablation Study for MolAug

In this section, we substitute MolAug with general graph augmentations including nodes drop (ND), edges perturbation (EP) in GraphCL [You *et al*., 2020] and domain knowledge-enriched molecular graph augmentations (DK) in MoCL [Sun *et al*., 2021]. Besides, we also consider chemical enantiomers (Enan.) and homologies (Homo.) of MolAug in isolation. The results demonstrated in Figure 5 illustrate that general graph augmentations and DK bring negative or limited improvements over vanilla fine-tuning, which may alter molecular property completely. Besides, using Enan. and Homo. in isolation is not the optimal alternative. Instead, the combinations of these two ingredients of MolAug achieve the best performance.

**Figure 5:**
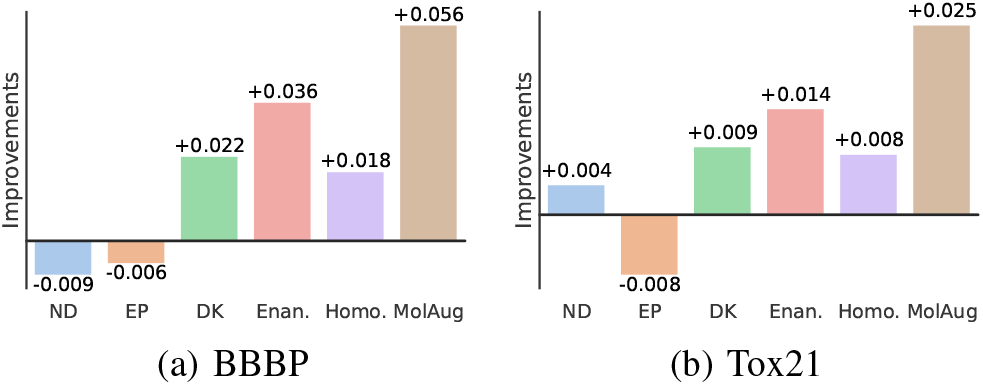
Comparisons between MolAug and existing molecular graph augmentations. The improvements refer to the improvements over vanilla fine-tuning.

#### Ablation Study for WordReg

We substitute the worst-case dropout with random dropout to study its influence. As shown in Figure 6, WordReg outperforms the random dropout across various drop ratios, which validates that looking for the worst-case dropout is necessary and effective. Figure 6(a) is symmetrical because both two dropouts are random.

**Figure 6:**
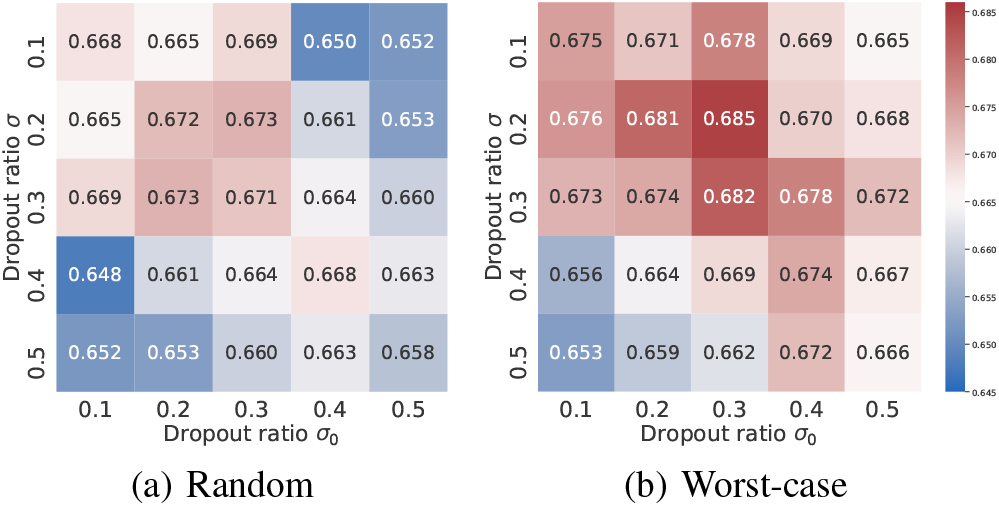
Comparisons between WordReg and random dropout regularization. The experiments are conducted with MPG base model on SIDER dataset. Here *σ*_0_ and *σ* are dropout ratio of vanilla training (fine-tuning) and worst-case drop, respectively.

### 4.6 Hyper-parameters Analysis

#### Worst-case dropout ratio

We have study dropout ratio in Figure 6(b). Obviously, WordReg can achieve brilliant performance when the two dropout ratios are in a reasonable range (0.1-0.3). Besides, WordReg tends to perform better when the two dropout ratios are close or identical.

#### Augmentations times of MolAug

As shown in Figure 7(a), more augmentations for each molecule at each iteration bring improvements. However, the performance is prone to be steady when the augmentations times are up to 3. On the other hand, more augmentations times will lead to unnecessary computational overhead.

**Figure 7:**
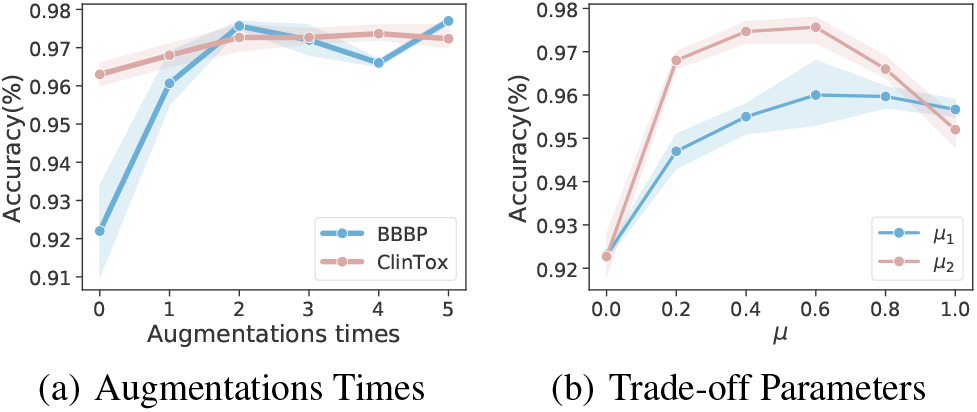
Hyper-parameters Analysis.

#### Trade-off parameters *μ*_1_, *μ*_2_

Figure 7(b) shows an different influence between *μ*_1_ and *μ*_2_. For *μ*_1_, the accuracy will be improved and then tend to converge with its value increasing, which illustrates that MolAug can also work well even if its trade-off parameter is oversized. In contrast, for *μ*_2_, the accuracy sees a dramatical drop, which can be imputed to its over-powerful regularization.

## 5 Conclusions and Future Work

We propose two effective strategies, MolAug & WordReg, for fine-tuning pre-trained molecular graph models to alleviate the over-fitting issues. The empirical results suggest that our strategies can advance the performance of vanilla fine-tuning on various pre-trained models and outperform state-of-the-arts results by a large margin. Also, we validate that our strategies are superior to existing fine-tuning techniques. Interesting direction for future work is applying our strategies to other molecule-related tasks such as molecule generation.

